# Aevol-9: A simulation platform to decipher the evolution of genome architecture

**DOI:** 10.1101/2025.04.10.648095

**Authors:** Juliette Luiselli, David P. Parsons, Romain Gallé, Paul Banse, Jonathan Rouzaud-Cornabas, Guillaume Beslon

## Abstract

Aevol is a forward-in-time simulator of genome architecture. It simulates a population of individuals, each with an explicit genome whose sequence and architecture can be modified by various mutational operators, including substitutions, indels, and large-scale chromosomal rearrangements. Aevol allows *in silico* experiments to decipher the effects of evolutionary conditions (*e*.*g*. population size, selection strength, mutation rates, and biases) on genome organization.

**Significance Statement:** The evolution of genome architecture is influenced by several factors, such as population size or mutation rates, but their impact is difficult to assess experimentally. We propose a simulation platform, named Aevol, that can be used to test *in silico* the effect of changes in several life traits on genome architecture. Aevol simulates a population of individuals, each with a structured double-stranded genome, that compete with one another and undergo various types of mutations, including chromosomal rearrangements that can affect genome architecture. This makes it possible to observe how genomes evolve in response to various selective and non-selective conditions. As such, Aevol is of great interest for any researcher wanting to study the evolution of genome architecture.

## Introduction

Genomes are multi-scale objects in which information is encoded at different levels: in the main sequence, but also in the composition of the genetic repertoire, which is a major source of variation (Chothia et al., 2003), and, at a higher level, in the genome architecture, defined as the non-random arrangement of the functional and non-functional elements in the genome (Koonin, 2009). Deciphering the laws of genome evolution hence requires understanding how these different levels evolve independently from one another and in conjunction with one another. Modeling and simulation have long proven to be decisive allies when it comes to understanding how genomes evolve under the combined action of selection, drift, and mutation (including substitutions, indels, chromosomal rearrangements, and recombination). Many simulation tools are available to explore evolution at the sequence level, often with an abundance of refinements. For example, one can cite SLiM (Haller et al., 2026), Nemo (Beeravolu Reddy et al., 2026) or AliSim (Ly-Trong et al., 2022). However, in these tools, genome architecture is defined by the user rather than free to evolve, even though its evolution remains poorly understood.

Genome architecture is very diverse between the different living kingdoms, ranging from very compact genomes in viroids and viruses (Gago et al., 2009) to very large – and apparently largely non-coding – genomes in some land plants (Pellicier et al., 2010), with prokaryotic genomes lying at intermediate complexity and architecture (Koonin, 2009). But genome architecture also seems to follow some hidden laws, as exemplified by the massive gene loss that is systematically observed in obligate endosymbionts or the strong genome streamlining observed in many marine bacteria (Batut et al., 2013). Such observations call for a better understanding of the laws governing the evolution of genome architecture and for the development of simulation tools to help understand how the macroscopic organization of functional and non-functional genomic elements responds to different evolutionary conditions (Mérot et al., 2020).

Aevol is a simulation platform specifically designed to study the evolution of genome architecture and gene repertoire. In Aevol, a population of individuals, each with a structured double-stranded genome, evolves under the combined action of selection, drift, and a wide range of mutations, including substitutions, indels, and chromosomal rearrangements. By varying conditions such as population size or the rates and biases of the different mutational events, one can observe how genome architecture responds, and relate the evolution of functional and non-functional genomic elements to the three main evolutionary forces: mutation, selection, and drift. Since its first use (Knibbe et al., 2007), Aevol has been continuously developed and applied to questions ranging from the fate of duplicated genes to the effects of population size and mutation rate on genome streamlining (Rutten et al., 2019; Liard et al., 2020; Banse et al., 2024b; Kalhor et al., 2024; Luiselli et al., 2024), but it has never been presented as a software platform in its own right.

Here, we describe Aevol in its current version, Aevol-9, which substantially broadens the range of questions the platform can address. In addition to the original prokaryote-like model (Standard), it includes a eukaryote-like version (Eukaryote), with linear diploid genomes, sexual reproduction, and recombination. A third version (4-Bases) uses a four-letter alphabet and the canonical genetic code, connecting simulated genomes to standard sequence analyses such as *dN/dS* estimation. Finally, its parallel implementation makes long experiments and large numbers of replicates computationally affordable. In the following, we first explain what makes the evolution of genome architecture difficult to simulate and how Aevol addresses this challenge, then describe the model, and finally illustrate its use through previous applications and a typical experiment.

## Why Aevol?

Any evolutionary simulator must link genetic variation to selection, that is, determine which mutations, alleles, or phenotypes become fixed in a population and which are eliminated by purifying selection. Most forward-in-time evolutionary simulators address this with one of two strategies. The first draws the effect of new mutations from a predefined distribution of fitness effects (DFE). This distribution can be refined by accounting for differences between coding and non-coding sequences, or for the position of a mutation within a reading frame. This strategy is implemented in widely used software such as SLiM (Haller et al., 2026) and Nemo (Beeravolu Reddy et al., 2026), and is well suited to investigating the fate of genetic variants under different population-genetic conditions. The second strategy relies on a predefined fitness landscape that maps genetic variants to fitness values. A recent example is STUN, which allows researchers to investigate how evolution proceeds on fitness landscapes with varying degrees of epistasis (Amado et al., 2023).

Neither of these strategies is well suited to simulating the evolution of genome architecture, primarily because of the complexity of the events that can modify genome organization. Genome architecture is shaped both by local-scale events, such as substitutions, small insertions and small deletions (collectively referred to as indels), and large-scale chromosomal rearrangements, such as inversions, large deletions, and segmental duplications. Incorporating chromosomal rearrangements into forward-in-time simulators is notoriously challenging for several reasons. First, their fitness effects remain poorly characterized, which makes it hard to specify an appropriate DFE in advance. Moreover, rearrangements can alter the DFE of subsequent mutations, so that the DFE itself changes over the course of evolution: gene duplication, for instance, can increase robustness to mutations affecting one of the two copies (Jouffrey et al., 2021). Second, these events can change genome size and gene copy number, which complicates the use of predefined fitness landscapes. Finally, rearrangements act at multiple scales: beyond altering gene repertoires through gene deletion or duplication, they can also generate novel coding sequences through gene fusion or fission, whose fitness effects are largely unknown.

For these reasons, most forward-in-time evolutionary simulators do not model chromosomal rearrangements at all. The few that include such events generally support a limited set of structural changes that move or invert genomic regions along a chromosome, without modeling their potential effects on gene sequences. Sequence evolution is then typically restricted to point mutations. However, there is no reason to assume that rearrangements occur exclusively between genes. Spontaneous intragenic rearrangements are also likely to occur, particularly in prokaryotes, where gene density is high. Although many of them probably disrupt gene function and thus are purged by purifying selection, they may nevertheless be a source of novel genetic variation (Chanin et al., 2024).

As a result, simulators capable of explicitly modeling the evolution of genome architecture are lacking. Aevol aims to fill this gap through an original way of assessing the fitness effects of genetic variation. Its central premise is that the fitness effects of genetic changes are largely determined by how they alter the encoding of genetic information. Aevol therefore computes each individual’s phenotype, and hence its fitness, directly from its genome, using an explicit, biologically inspired genotype-to-phenotype (G2P) map. Because any genome, however it was modified, can be decoded into a set of genes and then into a phenotype, Aevol can compute the fitness effect of virtually any genomic change, from local mutations to large structural rearrangements, including those that disrupt or fuse genes. Aevol thus follows an *in silico* experimental evolution approach (Hindré et al., 2012; Batut et al., 2013), in which substitutions, indels, structural variants, and their combinations can all be simulated over millions of generations. Throughout such experiments, one can track the fixation dynamics of mutations and the resulting changes in genomic features (*e*.*g*. the fraction of coding and non-coding sequence, the proportion of polycistronic mRNAs, or the mean number and size of genes), and thus investigate how each type of mutation contributes to the evolution of genome architecture.

Figure 1 provides an overview of the genotype-to-phenotype (G2P) mapping implemented in Aevol. The mapping comprises five main steps: transcription, ORF identification, translation and folding, metabolism, and fitness computation. These steps are inspired by the molecular mechanisms through which cells process information, particularly those described by the so-called “central dogma of molecular biology”. The first steps explicitly represent the decoding of genomic sequence, which is what allows genome architecture to evolve in the model. By contrast, the final steps rely on an abstract representation of biological functions. Although this abstraction departs from biological reality, it provides the computational efficiency required to simulate the evolution of genome architecture over several million generations.

**Figure 1.**
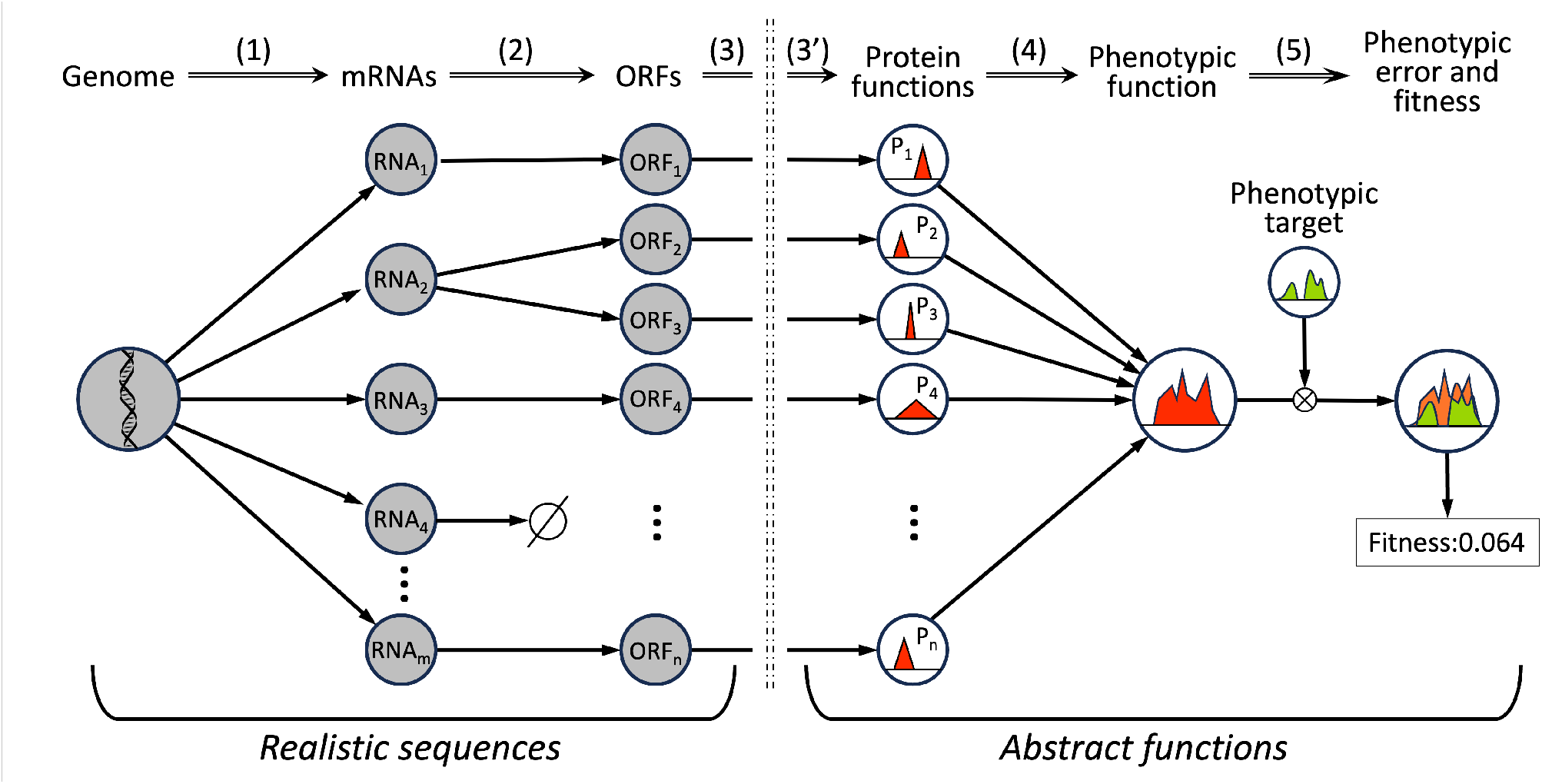
Overview of the Genotype-to-Phenotype map in Aevol. **(1) Transcription**. Both strands of the genome are parsed to identify mRNAs. Promoters are identified based on a predefined consensus sequence, and transcription proceeds along the DNA strand until a terminator (a sequence able to form a stem-loop) is encountered. The similarity between the promoter and the consensus sequence determines the transcription rate, *i*.*e*. the mRNA’s expression level *e*. **(2) ORF identification**. Each mRNA is scanned for translation initiation signals (a Shine-Dalgarno-like sequence followed by a start codon); the downstream sequence forms an Open Reading Frame (ORF) until a STOP codon is found in the same reading frame. A given mRNA may contain several ORFs (a polycistronic mRNA, or “operon”, *e*.*g*. RNA_2_) or none (a non-coding RNA, *e*.*g*. RNA_4_). **(3) Translation**. Each ORF is translated codon by codon using a predefined genetic code. In the Standard and Eukaryote versions of Aevol, an artificial binary genetic code is used, encoding six amino-acid (AA) types. In the 4-Bases version, the canonical genetic code is used. The first three steps use decoding processes that are directly inspired by the encoding of information in molecular biology, ensuring that any alteration of the genomic sequence will be propagated along the G2P map in a realistic way. The following steps move beyond sequence space and represent biological functions as mathematical functions in an abstract phenotypic space. **(3’) Folding**. The protein primary sequence is used to compute the function of the corresponding protein. Each AA sequence is decoded into three parameters defining a triangular function: *m* (position of the protein’s contribution along the phenotypic space), *w* (width, interpreted as pleiotropy), and *h* (height, *i*.*e*. activation or inhibition level). **(4) Metabolism**. Each protein’s contribution is weighted by its mRNA expression level *e*. Protein functions are then combined: activating and inhibiting functions are summed separately, and inhibition is then subtracted from activation. The resulting function represents the organism’s phenotype (red area). **(5) Fitness computation**. The phenotype is compared with a predefined phenotypic target function (green area). The integral of the absolute difference between the two functions (light red area) yields the phenotypic error *g*. Fitness is then computed as exp(*−kg*), where *k* controls the strength of selection.

The modeling choices made in Aevol sets it apart from other evolutionary simulators. Tools such as SLiM (Haller et al., 2026), Nemo (Beeravolu Reddy et al., 2026), or AliSim (Ly-Trong et al., 2022) offer great flexibility in the demographic, ecological, or phylogenetic scenarios they can simulate, but the genome architecture or the fitness effects of mutations are specified by the user. In Aevol, by contrast, genome architecture and the fitness effects of mutations are not inputs of the model but outcomes of evolution, emerging from the decoding of the genome. Aevol therefore does not aim to perform the same tasks as these tools, and a direct comparison would be of limited relevance. Instead, it provides an exploratory platform to test hypotheses about the evolution of genome architecture, for which no comprehensive theoretical framework currently exists, with the ultimate goal of contributing to the development of such frameworks.

## Technical description

A detailed description of the model is available on the website (https://www.aevol.fr/model-description/). In the following, we will only briefly present the main features of the model. Figure 2 illustrates the main characteristics of the model.

**Figure 2.**
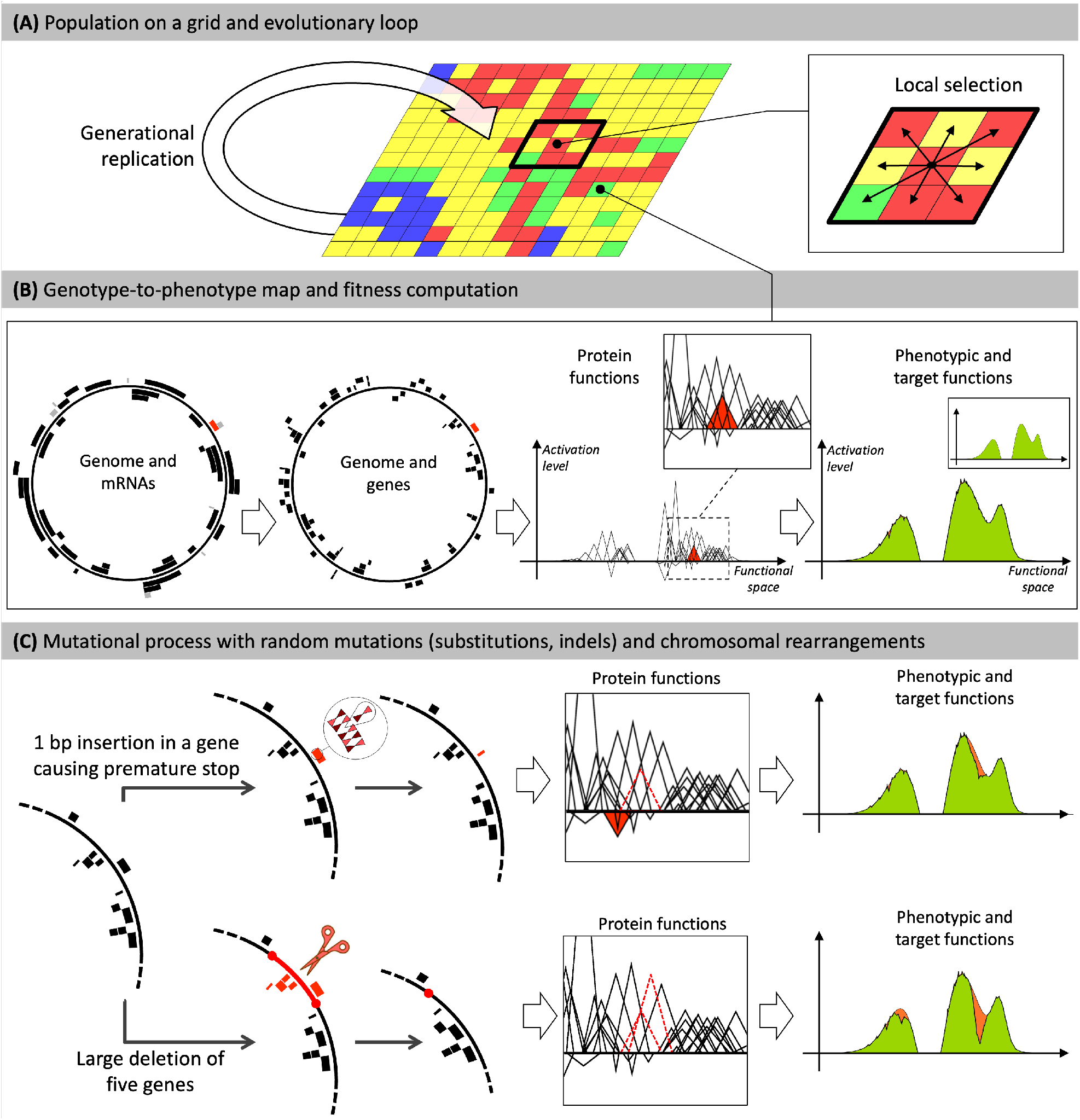
Overview of the Aevol model. **A) Population level**. Individuals are distributed on a grid. At each generation, the entire population replicates (generational model). Here, selection operates locally within a 3 × 3 neighborhood. **B)** Each grid cell contains a single organism, characterized by its genome. Genomes are decoded through a genotype-to-phenotype map comprising five main steps: transcription, translation, computation of protein functions, computation of the phenotype, and fitness computation. For illustration purposes, a given gene and its corresponding mRNA are shown in red. The red triangle represents the function of the corresponding protein in the model’s mathematical framework. Once the genome has been decoded, the phenotypic function is computed by summing the functions of all proteins. The phenotype is then compared with a predefined target (green area) to calculate fitness. The individual shown here evolved in the model during 500,000 generations in a binary, haploid setup. **C)** Individuals may undergo mutations during replication. Two examples are shown here: a small insertion (top) and a large deletion (bottom). **Top:** A 1-bp insertion within the red gene, causing a frameshift and introducing a premature stop codon. The gene’s ancestral function is lost (dashed triangle), and the resulting truncated protein has a different function (red triangle). This increases the difference between the phenotype and the target (light red area in the phenotype), resulting in a loss of fitness. **Bottom:** The deletion removes five genes. The functions of two of these genes are shown in the box (dashed triangles). The deletion results in a large discrepancy between the phenotype and the target (light red area in the phenotype). Figure reproduced from Luiselli et al. (2024).

### Population level

Aevol models a population of organisms on a grid that evolves through processes of variation, drift, and selection. The population follows a generational model (Fig. 2A): the whole population is replaced at each generation, and individuals compete to populate the next generation. The selection strength, population size, and grid structure can all be parameterized, and competition can be global or local, allowing users to investigate how drift and selection influence the evolution of genome architecture (Misevic et al., 2015; Luiselli et al., 2024). Reproduction is asexual in the haploid genome models and sexual, with meiotic recombination, in the diploid one (see below).

### Genome models and genotype-to-phenotype map

The phenotype of an individual is a mathematical function computed by decoding its genome through the G2P map described above (Fig. 1; Fig. 2B). Individuals are therefore fully characterized by the genetic information they carry. Aevol-9 offers three genome models: In the Standard version, genomes are haploid and circular, akin to prokaryotic genomes. In the Eukaryote version, genomes are linear and diploid. In both cases, the genetic sequences are represented by complementary binary nucleotides, translated with an artificial genetic code. By contrast, the 4-Bases version uses a 4 bases nucleotidic alphabet with a prokaryotic circular genome. In the 4-Bases version, translation relies on the canonical genetic code and its degeneracy can be used to estimate the mode of evolution (*dN/dS* ratio) or the synonymous and non-synonymous polymorphisms (*πN/πS* ratio).

### Mutational process

Upon reproduction, each individual can undergo sequence alteration through a set of variation operators: local mutations (substitutions, indels – see *e*.*g*. Fig. 2C, top, for an example of a small insertion of one base) or chromosomal rearrangements (deletions, inversions, duplications, and translocations – see *e*.*g*. Fig. 2C, bottom, for an example of a large deletion). In the diploid Eukaryote version, genomes can also undergo recombination events. Importantly, in Aevol, the selection coefficient of a new mutation is not drawn from a predefined distribution of fitness effects. Instead, it is computed by comparing the fitness of the parent with that of the mutant, after decoding the mutant genome. Consequently, the fitness effect of a mutation depends not only on the specific genomic change but also on the genetic background in which it occurs. This allows epistatic interactions between mutations to influence their fitness effects. The resulting DFE is therefore not fixed but can change over the course of evolution in response to changes in genomic architecture and genetic background. This makes it possible to investigate the evolutionary forces that shape the DFE.

### Implementation and performance

Aevol includes advanced parallelization, enabling fast simulations over the long evolutionary timescales required to observe the emergence of stable genome architectures. Its computational efficiency also makes it possible to perform a large number of independent replicates, thereby enabling robust statistical analyses. For instance, Luiselli et al. (2024) used populations pre-evolved for 10 million generations and performed 500 replicates in new conditions of population size, mutation rates, or both, for 2 million generations. Such large experimental campaigns are typically run on high-performance computing facilities, but smaller campaigns can be run on a laptop: the experiments presented in the following section were run on a MacBook Pro equipped with an Apple M3 Max chip, using 10-core parallel computation. The computational speed was approximately 400, 000 generations per hour for a population of 1024 individuals, scaling roughly inversely with population size. The entire experiment therefore took approximately three days to complete on this laptop.

## Use cases

### Emergent genome architectures

Because genome architecture is not specified in advance but results from the decoding of evolving sequences, a wide variety of genomic structures emerge spontaneously in Aevol simulations, including polycistronic mRNAs, non-coding RNAs, and overlapping or nested genes and mRNAs. These functional elements are interspersed with non-functional segments of variable size, whose accumulation can also be investigated. As a result, the same phenotypic information can be encoded by markedly different genome architectures, with different numbers of genes and coding densities.

### Typical workflow

Typical applications of Aevol involve running replicate simulations to determine whether specific genome architectures emerge consistently under given evolutionary conditions, such as population size, mutation rates and biases, mutation types, or reproductive regimes, and to relate these patterns to the underlying evolutionary forces. To support such investigations, Aevol provides tools for post-evolutionary analyses at the level of individuals, populations, or entire lineages. These include reconstructing the mutational history of an individual, including the selection coefficients of fixed mutations and rearrangements, estimating individual robustness and evolvability, and estimating the effective population size along an experiment. Combining these analyses makes it possible to disentangle the contributions of different parameters and evolutionary forces to the observed evolutionary trajectories.

### Previous applications

Aevol has been used to study how mutation rates and population size shape genome architecture, from the amount of non-coding DNA (Knibbe et al., 2007) to the effect of mutator alleles (Rutten et al., 2019) and the conditions for genome streamlining (Luiselli et al., 2024). Other studies have focused on genes and the events that create or remove them: the evolution of gene repertoire complexity (Liard et al., 2020), the fate of duplicated genes (Kalhor et al., 2024), and the role of chromosomal rearrangements (Banse et al., 2024b), showing for instance that segmental duplications can cause evolutionary bursts in viruses, even in constant conditions (Banse et al., 2024a). Finally, because the complete mutational history of simulated genomes is known, Aevol can also serve as a benchmark for bioinformatic tools (Biller et al., 2016; Daudey et al., 2024).

### An illustrative experiment

To illustrate the full workflow, we present a simple experiment investigating the effects of population size on genome architecture. The role of genetic drift in shaping genome size remains debated. The mutational-hazard hypothesis (MHH) proposes that differences in genome size and architecture across species can be explained, at least in part, by the interplay between mutations and genetic drift (Lynch and Conery, 2003), but recent comparative genomic analyses have questioned its generality (Marino et al., 2025). Aevol makes it possible to test predictions of the MHH through controlled *in silico* experiments.

Starting from a bacteria-like strain that had evolved in Aevol (Standard version) for 10 million generations in a constant-size population (*N* = 1024 individuals), we used `aevol_create_2b` to initiate 15 simulations with three different population sizes (*N*, *N/*4, and 4*N*, with five replicates each), while keeping all other conditions identical to those used during the wild-type evolution. Specifically, we used the `WT2_bact_2b` strain, available on the Aevol website (https://www.aevol.fr/wildtypes/), which has the median genome size among the five bacteria-like wild-types available on the website. We then used `aevol_run_2b` to let these 15 populations evolve for 1.1 million generations. After the simulations were finished, the lineage leading to the final best individual was recovered using `aevol_post_lineage_2b`, and the genomic characteristics of the ancestral lineage were reconstructed using `aevol_post_ancestor_stats_2b` for subsequent analysis.

Figure 3 shows the main results of this experiment. We show individual trajectories rather than averages, to provide a visual representation of the evolutionary dynamics generated by the model. Fitness (top left) reflects how population size affects the balance between mutation, selection, and genetic drift. Here, the impact of population size on the balance is clearly reflected in the magnitude of fitness fluctuations, which arise from differences in the fixation probabilities of deleterious mutations. Specifically, reducing population size shifts the balance towards drift, resulting in a decline in fitness (blue curves). By contrast, when population size is maintained (red curves) or increased (green curves), fitness tends to increase gradually, with occasional large jumps. These stepwise fitness gains illustrate the punctuated pattern of evolution generated by interactions between different types of mutational events (Banse et al., 2024a), with larger populations exhibiting more frequent jumps. In the top right panel, each simulation is represented by two curves: the lower curves show changes in coding genome size across generations (with all simulations overlapping at this scale, indicating that coding DNA content is only marginally affected in this experiment), while the upper curves show changes in total genome size (*i*.*e*., the combined coding and non-coding compartments). This representation allows to visualize changes in the total amount of DNA (upper curve), in coding DNA content (below the lower curve) and non-coding DNA content (between the two curves). Increasing population size leads to a reduction in total genome size, primarily driven by a contraction of the non-coding compartment. This is consistent with the mutational-hazard hypothesis (MHH), while also raising interesting questions about the mechanisms underlying genome size variation. By contrast, changes in coding DNA appears to be limited and largely independent of population size (inset).

**Figure 3.**
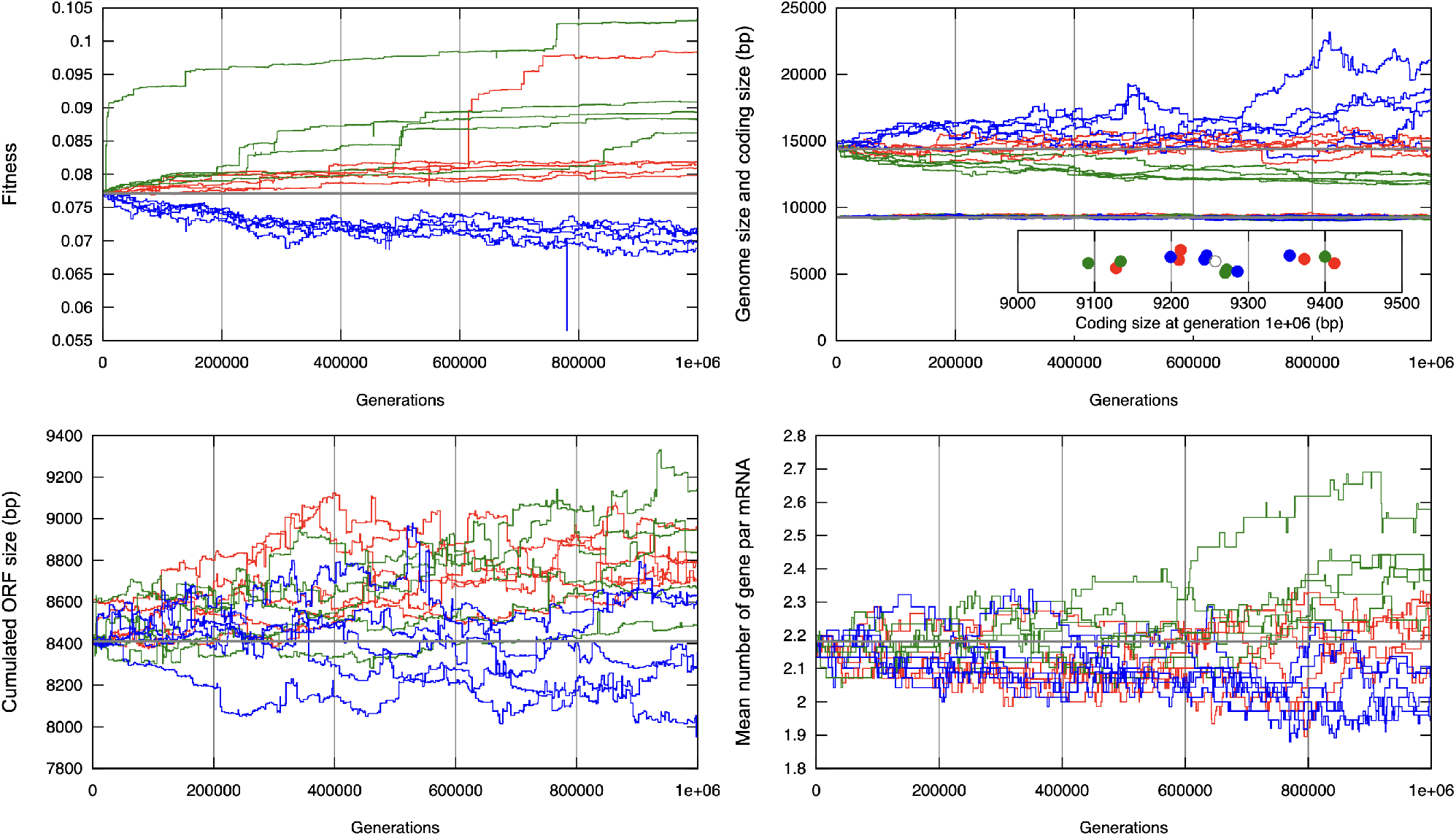
A typical experiment with Aevol. Starting from a pre-evolved wild-type strain, 15 independent simulations were run with 3 population sizes: *N* = 1024 (red; as during wild-type evolution), 4*N* = 4096 (green), and *N/*4 = 256 (blue). The four panels show evolutionary dynamics along the lineage leading to the best individual at generation 1.1 *×* 10^6^. The last 100, 000 generations were excluded to ensure that all reported events had reached fixation. In all panels, gray horizontal lines indicate generation-0 values (also marked by a gray circle in the inset of the top-right panel). **Top left**: fitness. **Top right**: genome size (top curves) and coding size (bottom curves, all simulations effectively overlapping at that scale); the difference represents non-coding genome size. The inset zooms in on coding size at generation 10^6^. **Bottom left**: cumulative length of genomic ORFs. **Bottom right**: mean number of ORFs per mRNA (mean operon size). See the main text for discussion.

The two bottom panels provide further insight into the organization of the coding compartment. They show that, although the amount of coding DNA does not depend on population size in our experiments (inset in the top-right panel), the cumulative length of ORFs does. This suggests that fitness losses in small populations and fitness gains in large populations are associated with decreases and increases, respectively, in the amount of genetic information. At the level of the coding compartment as a whole, however, these changes are compensated by changes in how the information is organized: genome size and operon size are inversely related, shorter genomes tending to have more ORFs per mRNA (bottom right). These results suggest that genome architecture is shaped by complex mechanisms involving both selective and non-selective forces. Although a detailed analysis of these dynamics is beyond the scope of this article, they illustrate the potential of Aevol as an experimental framework for investigating the evolution of genome architecture.

## Conclusion

Aevol enables unique evolutionary experiments to study the evolution of genome architecture and its dependence on many parameters. In that respect, it meets a need that, to our knowledge, no other model currently fulfills. Recent developments allow large-scale and long-lasting experiments with unprecedentedly strong statistical power, and new versions of Aevol (4-Bases and Eukaryote) open a whole new range of possibilities to decipher the different evolutionary forces acting on genome architecture. Furthermore, the model can be easily extended to explore new avenues like the evolution of gene regulation networks, the evolution of polyploidy, or the role of genetic dominance on the fate of duplicated genes.

## Implementation and data availability

Aevol is available for download at https://www.aevol.fr/doc/download/. Documentation and examples to demonstrate usage and possible experiments are available at (www.aevol.fr/doc/). Aevol is implemented in C++20 and uses OpenMP. It is compatible with Linux and MacOS, provided the C++20 compilation toolchain is available. Data of the simulations presented in this paper are available on Zenodo: https://doi.org/10.5281/zenodo.24009273.

## Author contributions

J.L, D.P, R.G., J.R.C, and G.B. contributed to the development of the software. All authors contributed to discussions about its usage and the manuscript. All authors wrote and reviewed the manuscript and the documentation of the software.

Conflict of interest: None declared.

## Bibliography

Amado, A., Li, J., and Bank, C. (2023). Stun: forward-time simulation on tunable fitness landscapes in recombining populations. Bioinformatics advances, 3(1):vbad164.

Banse, P., Elena, S. F., and Beslon, G. (2024). Innovation in viruses: fitness valley crossing, neutral landscapes, or just duplications? Virus Evolution, 10(1):veae078. doi: 10.1093/ve/veae078.

Banse, P., Luiselli, J., Parsons, D. P., Grohens, T., Foley, M., Trujillo, L., Rouzaud-Cornabas, J., Knibbe, C., and Beslon, G. (2024). Forward-in-time simulation of chromosomal rearrangements: The invisible backbone that sustains long-term adaptation. Molecular Ecology, 33(24):e17234. doi: 10.1111/mec.17234.

Batut, B., Parsons, D. P., Fischer, S., Beslon, G., and Knibbe, C. In silico experimental evolution: a tool to test evolutionary scenarios. In BMC bioinformatics, volume 14, pages 1–11. Springer, (2013).

Beeravolu Reddy, C., Chebib, J., Cotto, O., Schmid, M., and Guillaume, F. (2026). Nemo2. 4: fast and accurate quantitative genetics forward-time simulations. bioRxiv, pages 2026–07.

Biller, P., Knibbe, C., Beslon, G., and Tannier, E. Comparative genomics on artificial life. In 12th Conference on Computability in Europe, Paris (France), July 2016, pages 35–44. Springer, (2016).

Chanin, R. B., West, P. T., Wirbel, J., Gill, M. O., Green, G. Z., Park, R. M., Enright, N., Miklos, A. M., Hickey, A. S., Brooks, E. F., et al. (2024). Intragenic dna inversions expand bacterial coding capacity. Nature, 634(8032):234–242.

Chothia, C., Gough, J., Vogel, C., and Teichmann, S. A. (2003). Evolution of the protein repertoire. Science, 300(5626):1701–1703.

Daudey, H., Parsons, D. P., Tannier, E., Daubin, V., Boussau, B., Liard, V., Gallé, R., Rouzaud-Cornabas, J., and Beslon, G. Aevol_4b: Bridging the gap between artificial life and bioinformatics. In ALIFE 2024: Proceedings of the 2024 Artificial Life Conference. MIT Press, (2024).

Gago, S., Elena, S. F., Flores, R., and Sanjuán, R. (2009). Extremely high mutation rate of a hammerhead viroid. Science, 323 (5919):1308–1308.

Haller, B. C., Ralph, P. L., and Messer, P. W. (2026). Slim 5: Eco-evolutionary simulations across multiple chromosomes and full genomes. Molecular Biology and Evolution, 43(1):msaf313.

Hindré, T., Knibbe, C., Beslon, G., and Schneider, D. (2012). New insights into bacterial adaptation through in vivo and in silico experimental evolution. Nature Reviews Microbiology, 10(5):352–365.

Jouffrey, V., Leonard, A., and Ahnert, S. (2021). Gene duplication and subsequent diversification strongly affect phenotypic evolvability and robustness. Royal Society Open Science, 8(6):201636.

Kalhor, R., Beslon, G., Lafond, M., and Scornavacca, C. (2024). A rigorous framework to classify the postduplication fate of paralogous genes. Journal of Computational Biology, 31(9):815–833. doi: 10.1089/cmb.2023.0331.

Knibbe, C., Coulon, A., Mazet, O., Fayard, J.-M., and Beslon, G. (2007). A long-term evolutionary pressure on the amount of noncoding dna. Molecular biology and evolution, 24(10):2344–2353.

Koonin, E. V. (2009). Evolution of genome architecture. The international journal of biochemistry & cell biology, 41(2):298–306.

Liard, V., Parsons, D. P., Rouzaud-Cornabas, J., and Beslon, G. (2020). The complexity ratchet: Stronger than selection, stronger than evolvability, weaker than robustness. Artificial life, 26(1):38–57.

Luiselli, J., Rouzaud-Cornabas, J., Lartillot, N., and Beslon, G. (2024). Genome streamlining: Effect of mutation rate and population size on genome size reduction. Genome Biology and Evolution, 16(12):evae250. doi: 10.1093/gbe/evae250.

Ly-Trong, N., Naser-Khdour, S., Lanfear, R., and Minh, B. Q. (2022). Alisim: a fast and versatile phylogenetic sequence simulator for the genomic era. Molecular biology and evolution, 39(5):msac092.

Lynch, M. and Conery, J. S. (2003). The origins of genome complexity. science, 302(5649):1401–1404.

Marino, A., Debaecker, G., Fiston-Lavier, A.-S., Haudry, A., and Nabholz, B. (2025). Effective population size does not explain long-term variation in genome size and transposable element content in animals. Elife, 13:RP100574.

Mérot, C., Oomen, R. A., Tigano, A., and Wellenreuther, M. (2020). A roadmap for understanding the evolutionary significance of structural genomic variation. Trends in Ecology & Evolution, 35(7):561–572.

Misevic, D., Frénoy, A., Lindner, A., and Taddei, F. (2015). Shape matters: Lifecycle of cooperative patches promotes cooperation in bulky populations. Evolution, 69. doi: 10.1111/evo.12616.

Pellicier, J., Fay, M. F., and Leitch, I. J. (2010). The largest eukaryotic genome of them all? Botanical Journal of the Linnean Society, 164(1):10–15. doi: 10.1111/j.1095-8339.2010.01072.x.

Rutten, J. P., Hogeweg, P., and Beslon, G. (2019). Adapting the engine to the fuel: mutator populations can reduce the mutational load by reorganizing their genome structure. BMC evolutionary biology, 19:1–17.

